# Gene-by-Environment Interaction Significantly Drives Bacterial Endophyte Communities in Maize Stalks

**DOI:** 10.64898/2025.12.23.696053

**Authors:** Hanxia Li, Holly Kristina Griffis, Jode Edwards, Sherry Flint-Garcia, Candice N. Hirsch, James B. Holland, Joseph E. Knoll, Dayane Cristina Lima, Natalia de Leon, Seth C. Murray, Maninder P. Singh, James C. Schnable, Rajandeep S. Sekhon, Peter Thomison, Addie Thompson, Jason G. Wallace

## Abstract

1.

**Background:** Many microbes are known to boost the performance of crop plants, and their use as crop treatments (“biologicals”) is an active area of research and commercialization. Although many identified microbes can boost plant production under laboratory settings, most fail to translate to field settings, and even the ones that are commercialized are often unreliable. While many factors likely underlie these issues, one that has been underexplored is the idea that interactions between plants and microbes can vary across both different host genetics and different environments, and that specific combinations of host and environment can result in very different microbial outcomes.

**Results:** To quantify how a plant microbiome can change across host genotype and environment and to determine the relative importance of these factors, we sampled maize stalks from 20 specific hybrid varieties grown at 15 locations in the United States. Using 16S rRNA gene sequencing, we find that the stalk microbiome is highly variable, with a handful of bacterial taxa conserved at broader taxonomic levels (Phylum, Class) and almost none at finer levels (Genus, Species). Local environment and gene-by-environment interactions both had consistently large and significant effects on the microbial community (measured by alpha-and beta-diversity summary statistics, individual taxa, and inferred functional capacity), whereas host genotype had little to no consistent effect across environments. We also identified specific soil factors (pH and potassium) as the most significant environmental drivers of the community, even though these communities were living inside the stalks.

**Conclusions:** Taken together, our results indicate that while host genetics can significantly affect the stalk microbial community, these effects are almost entirely predicated on interactions with the environment. These results imply that reliable deployment of microbes for crop production will require significant investment in trials across many locations and genetic backgrounds, either to estimate the GxE effects directly or to screen for microbes whose effects are less dependent on these two factors.

## 2. Background

In recent years, the scientific community has intensively investigated plant-microbe interactions for benefits in plant growth and production. For example, arbuscular mycorrhizal fungi, which are obligate symbionts, facilitate mineral and water uptake of plant roots [1]. Some rhizobacteria, such as *Pseudomonas spp*., have been found to suppress phytopathogens, stimulate plant growth by nitrogen fixation, and solubilize inorganic minerals [2–4]. Many such microbes have been isolated from plants (both cultivated and wild) for growth testing in the laboratory before being tested in the greenhouse and field. Collectively, these crop treatments are called “biologicals”, with an estimated market value of US$17 billion in 2025 and expected to grow to over $44 billion by 2032 [5].

However, growth-promoting microbes that perform well in the laboratory and under greenhouse conditions often fail when deployed as external inputs in real-world agricultural production [6]. Several factors are likely involved in this failure to translate products, including increasing environmental variability, poor adaptation to field conditions, variable interactions with different host genotypes, and difficulties with product formulation, among other unknowns to be discovered. Particularly, it has been shown that both the genotype of the host plant and the local environment affect the degree of benefit from specific microbes. For example, the growth promotion effect of *Azospirillum spp*. applied to sugarcane was found to be influenced by both drought stress and different plant genotypes [7], and different genotypes of common beans show contrasting responsiveness to the inoculation of *Azospirillum brasilense* [8]. Similarly, *Herbaspirillum seropediceae*, a nitrogen-fixing endophyte, promotes growth when applied to only some genotypes of maize [9].

Geneticists and breeders have commonly observed that different plant genotypes will respond differently to environmental factors at measured phenotypes [10,11]. This “genotype-by-environment interaction” explains why one variety outperforms another in one environment (for, e.g., grain yield), but the relative values change in a different environment. Since plant-associated microbes respond to host metabolism [12], it is reasonable to assume that genotype-by-environment interaction within the host will also influence its interactions with microbial communities. However, the degree to which host genetics, environment, and their interaction alters plant microbiomes has received relatively little study, especially for major crops like maize. For example, Wagner et al [13] surveyed the leaf and root microbiomes of *Boechera stricta*, a perennial wild mustard, and found that host genetics affected leaf bacterial communities but not root ones, and that the degree of this effect varied significantly across environments. Edwards *et al.* [14] meanwhile, found significant genetic and environmental effects on the switchgrass (*Panicum virgatum*) root microbiome, with locally adapted genotypes preferentially recruiting local bacteria from their native range. A study of *Arabidopsis* at 267 locations across its native range found both environment (latitude, drought) and host genotype to significantly impact which bacteria were living on its leaves [15]. Finally, a prior study on the maize rhizosphere found that the environment was the largest influence on the bacterial community, whereas gene-by-environment interaction was a smaller but significant influence and host genetics by itself explained very little of the variation [16].

Since 2014, the U.S. maize community has been running large, public experiments on genotype-by-environment interaction through the Genomes to Fields (G2F) Initiative [17]. G2F consists of a grassroots collaboration among 25-30 field locations each year, each of which contains two replicates of at least 250 unique maize hybrids, with the specific hybrids changing every two years. This project ranks among the largest public datasets for agriculture available, and has been used to tackle a diversity of questions including using machine learning to predict plant performance [18,19], extracting disease and performance data from unmanned aerial vehicle imaging [20,21], recalibrate crop growth models [22], and investigate gene-by-environment interactions [17,23].

For this study, we sampled maize stalks across G2F locations and used 16S rRNA gene sequencing to survey the bacterial microbiome present in maize stalks. Our primary goals were to identify the nature of the maize stalk community; determine how much it varied across large geographic distances; and determine the relative effect of host genetics, environment, and GxE on the community makeup.

## 3. Methods

### 3.1 Maize stalk sampling

All sampled maize plants were part of the 2019 G2F experiments [24]; agronomic and phenotypic data for these experiments were published separately [25]. In brief, we selected 20 maize hybrid genotypes to sample from among the controls planted at each location. Pre-labeled collection bags were distributed to each collaborating location. Locations included Delaware, Georgia (2 sites), Iowa (2 sites), Indiana, Michigan, Minnesota, Missouri, North Carolina, Nebraska (2 sites), New York (2 sites), Ohio, South Carolina, and Texas (two sites), for 18 total locations. Sampling was performed slightly before peak flowering at each location, as determined by the local collaborator. For each plot, a random plant in the middle of a row was selected, and a 10-cm stalk segment was cut from 20 to 30 cm above the ground. Any leaves around the segment were removed, and the 10-cm stalk sample was put into a ziplock bag, which was then stored in a cooler filled with ice and ice packs. Clippers were cleaned with alcohol between each sample to remove carryover. (A video detailing this procedure was given to each collaborator and is available online [26].) After collection, all samples were shipped on ice to the University of Georgia via overnight shipping. The samples were stored in a 4°C refrigerator until DNA extraction (<48 hours later). In total, we received 840 stalk samples across all locations. An additional sampling occurred in 2020; however, most samples showed signs of exogenous bacterial contamination from an unknown source so only the 2019 data are included in the current analysis.

During sample processing, it was discovered that shortages in some seed stocks had resulted in last-minute seed substitutions, so that some samples came from different genotypes than planned for. Therefore, in downstream analyses a “corrected pedigree” column was added to the metadata to indicate the actual genotype sampled, which was used for the genotype data in all subsequent analyses. A small number of samples where it was unclear if the substitution had occurred—indicated by a blank “pedigree” field in the final G2F dataset—were given unique identifiers “Unknown1”, “Unknown2”, etc., so they could be used in environmental comparisons but would be filtered out of any genotype-specific analyses.

### 3.2 DNA extraction and sequencing

Samples were extracted by using a fresh razor blade to cut off 1-2 cm at each end of the stalk sample, which was then discarded. For the first three locations (TXH1, TXH2, GAH1), the outside of the stalk was removed with a fresh razor blade, and the interior was chopped. All later samples instead used a sterile 3/8-inch drill bit to extract the pith of the stalk, which was collected in a fresh weigh boat. Both the cutting boards and drill bits were cleaned with water and 70% ethanol, then UV-sterilized for 30 minutes between samples. The later downstream analysis found that the samples from the first three locations contained a high percentage of plastid sequences and were therefore excluded from the later analysis. The pith was then transferred into a 2mL tube containing bashing beads. Stalks were processed in batches of 18 (the number of drill bits available) and then immediately extracted with a Quick-DNA Fecal/Soil Microbe 96 Kit (Zymo), following the manufacturer’s instructions. Extracted DNA was stored at −20° C until all samples were received and processed.

16S rRNA gene amplification was performed using the Earth Microbiome Project 515F (Parada) and 806R (Apprill) primers [27], with Illumina barcode linkers and staggered spacers added [28]. Primer sequences are TCGTCGGCAGCGTCAGATGTGTATAAGAGACAG-XXX-GA<u>GTGYCAGCMGCCGCGGTAA</u> (515F) and GTCTCGTGGGCTCGGAGATGTGTATAAGAGACAG-XXX-GA<u>GGACTACNVGGGTWTCTAAT</u> (806R), where the first segment is the Illumina linker, “-XXX-” is a 0-5 bp staggered spacer [28], followed by a GA linker and the primer sequence itself (underlines). Commercial peptide nucleic acids pPNA and mPNA (PNA Bio), to block plastid and mitochondrial amplification, respectively, were mixed and diluted to 2.5µM each for inclusion in the reaction. The first PCR reaction consisted of 5 µL DNA template, 2 µL of each primer (0.5 µM), 12.5 µL of Hot Start Taq 2X Master Mix (New England Biolabs), 2.5 µL PNA mixture (2.5 µM each), and 1uL of sterile water. The amplification reaction was 95°C for 45 seconds; twenty cycles of 95°C for 15 seconds, 78°C for 10 seconds, 60°C for 45 seconds, 72°C for 45 seconds; and finally, held at 4°C. PCR products were purified with magnetic AMPure beads (Beckman Coulter Life Sciences).

Five µL of the first PCR product for each sample was used in the second PCR amplification to add sequencing barcodes and adaptors. The reaction mixture consisted of 5 µL first PCR product, 5 µL Nextera i5 plus i7 Barcode Primers (Illumina), 12.5 µL 2x Taq DNA polymerase master mix, and 2.5 µL PNA mix. The second amplification reaction was 95°C for 45 seconds; 25 cycles of 95°C for 15 seconds, 78°C for 10 seconds, 60°C for 45 seconds, 72°C for 45 seconds; and finally, 68°C for 5 minutes, followed by the hold at 4°C. The second PCR products were purified using AMPure beads and eluted in 27uL of sterile water. 25 µL of the solution from each sample, which was the final 16S rRNA gene sequencing library, was stored in the freezer at −20°C until sequencing. Three blanks were used in DNA extraction and library preparation. Libraries were sequenced at the Georgia Genomics and Bioinformatics Core on an Illumina MiSeq instrument using one paired-end 250 bp flowcell. Raw sequencing data are available in the NCBI SRA database (accession PRJNA883673).

### 3.3 Data preprocessing and quality control

Processing of raw sequencing data to amplicon sequence variants (ASVs) was done using the QIIME2 pipeline [29]. In brief, Cutadapt [30] was used to remove adapters from raw pair-end reads. The trimmed reads were imported into Qiime2 and clipped to 240 bp to avoid low-quality regions near the tail of the reads. Denoising and ASV calling were done with the DADA2 algorithm implemented in the Qiime 2 plugin [31]. Taxonomy for each ASV was assigned using a prebuilt 515F/806R region classifier based on the Silva v138 99% ASVs dataset [32] and supplied on the QIIME2 data resources page [33]. After the assignment, any mitochondria and chloroplast sequences were filtered out. The remaining ASV table contained 602 samples and 19,241 ASVs.

### 3.4 Data Joining

The sample metadata (including corrected pedigree names; see section 3.1), ASV table, assigned taxonomy, and phylogenetic tree from QIIME2 were imported into R and combined into a Phyloseq object [34]. ASVs present in blank samples were removed and equivalent genotypes were standardized (e.g., genotypes OH43/B73 and B73/OH43, which differ only in which plant was used as pollen or seed parent). Based on manual inspection of the data, all samples were rarefied to a read depth of 1,500, which left 502 samples and 17,909 ASVs.

### 3.5 Core Microbiome Analysis

To identify core bacterial taxa in the stalk microbiome, we first combined ASVs at major taxonomic levels (Phylum, Class, Order, Family, Genus, and Species) using the tax_glom() function in phyloseq v1.42.0 [34]. Core taxa were then defined as any taxon present at a relative abundance of 0.1% or more in at least 60% of samples. Using these criteria, we identified core microbes for the overall dataset, each location (15 total), and each maize genotype with at least 10 samples (19 total). Taxonomic trees were visualized with metacoder v0.3.8 [35] and manually adjusted in Inkscape [36].

### 3.6 Alpha and Beta Diversity Analysis

Alpha diversity metrics (Shannon index, Simpson index, and the number of observed features) were estimated using the estimate richness() function in phyloseq v1.42.0 [34], using groups (locations or genotypes) with at least 10 samples each. Statistical differences among groups were tested via the Kruskal-Wallis test. Since the UniFrac ordination method in Phyloseq has been deprecated, the beta diversity metrics (Weighted and Unweighted) were calculated with the beta.div() function in the package rbiom [37].

### 3.7 Quantifying genetic and environmental effects on microbiome composition

Each microbiome metric (alpha diversity, beta diversity, taxonomy, and inferred functionality [see below]) was analyzed for the relative contribution of genetics, environment, GxE, and unexplained residual error. Since some genotypes were only present in a single environment or only once in a given environment, we filtered the rarefied sample set down to those genotypes present at least twice per location in at least 3 locations, and to locations with at least 10 quality samples each. This resulted in a dataset of 323 samples across 10 locations (the “GxE subset”), containing 17,909 ASVs.

#### 3.7.1 Alpha diversity

Prior to analysis, the Observed ASVs metric was log-transformed and the Simpson index was reflected (subtracted from 1) and then log-transformed to improve normality. For all metrics, up to 10 outliers were removed with the Rosner’s test for multiple outliers [38] as implemented in the EnvStats package for R [39]. Ordinary least squares regression was then used to test the effect of environment, maize genotype, and their interaction with the equation

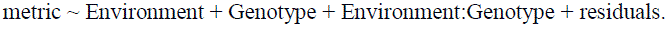

The resulting sum-of-squares for the environment, maize genotype, and GxE was calculated by type I ANOVA. The variation explained was estimated by dividing each term’s sum of squares by the total (including residuals).

#### 3.7.2 Beta diversity

For beta diversity, the Jaccard, Bray-Curtis, weighted UniFrac, and unweighted UniFract distance metrics were re-calculated for just the GxE subset of samples. The fraction of variation explained was calculated by distance-based redundancy analysis, as implemented in the dbrda() function in the vegan package for R [40], with ANOVA results determined from 999 permutations.

#### 3.7.3 Taxonomy

For individual taxa, the ASV table was collapsed at different taxonomic levels (phylum, class, order, etc.) and filtered to taxa with at least 20% prevalence. (That is, present in at least 20% of samples.) The variation explained was then estimated as per alpha diversity.

#### 3.7.4 Inferred functionality

The PICRUST2 pipeline [41] was used to estimate metabolic pathway abundances by MetaCyc ortholog based on the ASV table from the GxE subset, and pathways missing in more than 20% of samples were removed. For each inferred pathway, the data was transformed by subtracting the minimum value and then adding 1. (Since the next step involves a power transformation, values of 0 could result in undefined results.) Box-Cox transformation [42] was then performed, testing values of lambda between −2 and +2 and selecting the optimal value for each pathway separately. The transformed data was then analyzed by ordinary least squares and ANOVA, as per alpha diversity.

### 3.8 Correlation between microbiome diversity and environmental factors

Soil data for 2019 from the G2F maize project [25] were used for correlation analysis. Soil factors included soil pH, soluble salts in millimhos per centimeter (mmho cm^−1^), Organic Matter (Loss on Ignition method; LOI %), Nitrate (µg g^−1^), Potassium (µg g^−1^), Sulfate (µg g^−1^), Calcium (µg g^−1^), Magnesium (µg g^−1^), Sodium (µg g^−1^), Cation exchange capacity (CEC)/Sum of Cations (me 100g^−1^), %H Saturation, %K Saturation, %Ca Saturation, %Mg Saturation, %Na Saturation, Mehlich P-III (extractable phosphate) (µg g^−1^), % Sand, % Silt.

For alpha diversity, a series of linear regression models were fit that tested each soil parameter individually for association with median alpha diversity at that location. (One location in North Carolina, NCH1, did not record soil data in 2019 and so was excluded from this analysis.) For beta diversity, the rarefied reads of each sample were grouped by location and re-rarefied to a total read depth of 18,000 (the minimum of any location). The beta diversity between each location was estimated by Bray-Curtis dissimilarity and by weighted and unweighted UniFrac distances, while environmental distances were calculated by Euclidean distance. The relationship between beta diversity distance and environmental factor distance was then determined by a Mantel test with 999 permutations in the vegan package for R [40].

For inferred functionality, the predicted amount of each pathway was averaged for each location, then associated with soil parameters using MaAsLin2 [43].

### 3.8 Software Packages

In addition to software mentioned earlier, this analysis made use of the following software packages: argparse [44], data.table [45], dplyr [46], EnvStats [47], ggnewscale [48], ggpmisc [49], ggpubr [50], gridExtra [51], gvlma [52], MASS [53], pals [54], qiime2R [55], readr [56], Rtsne [57], stringr [58], tibble [59], and tidyverse [60].

## 4. Results

Maize stalks were sampled from control genotypes included in the 2019 Genomes to Fields experiments [24]. Each sample consisted of a 10-cm segment cut near the ground (20-30 cm from soil) just before flowering, which was kept on ice until extraction. (See Methods for details.) In total, we collected 843 samples across 18 locations.

After filtering out low-quality and organelle sequences, the amplicon sequence variant (ASV) table contained 602 samples 19,241 ASVs, with 2,265,108 high-quality reads and a median sequence depth of 3309 per sample. A rarefaction depth of 1,500 reads was selected to avoid the influence of different library sizes of samples on the microbiome composition, and any sample that did not have at least 1,500 reads was discarded. In the end, the rarefied ASV table kept 17,909 ASVs in 502 samples across 15 locations (**Figure 1, Supplementary Tables S1 & S2**). We had originally targeted the same 20 genotypes at each location, but last-minute seed substitutions meant that we actually captured 99 unique genotypes (including 20 “unknown” samples). However, eighty percent of samples (399 of 502) still belonged to the 19 most abundant genotypes, each of which was sampled at least 10 times.

**Figure 1:**
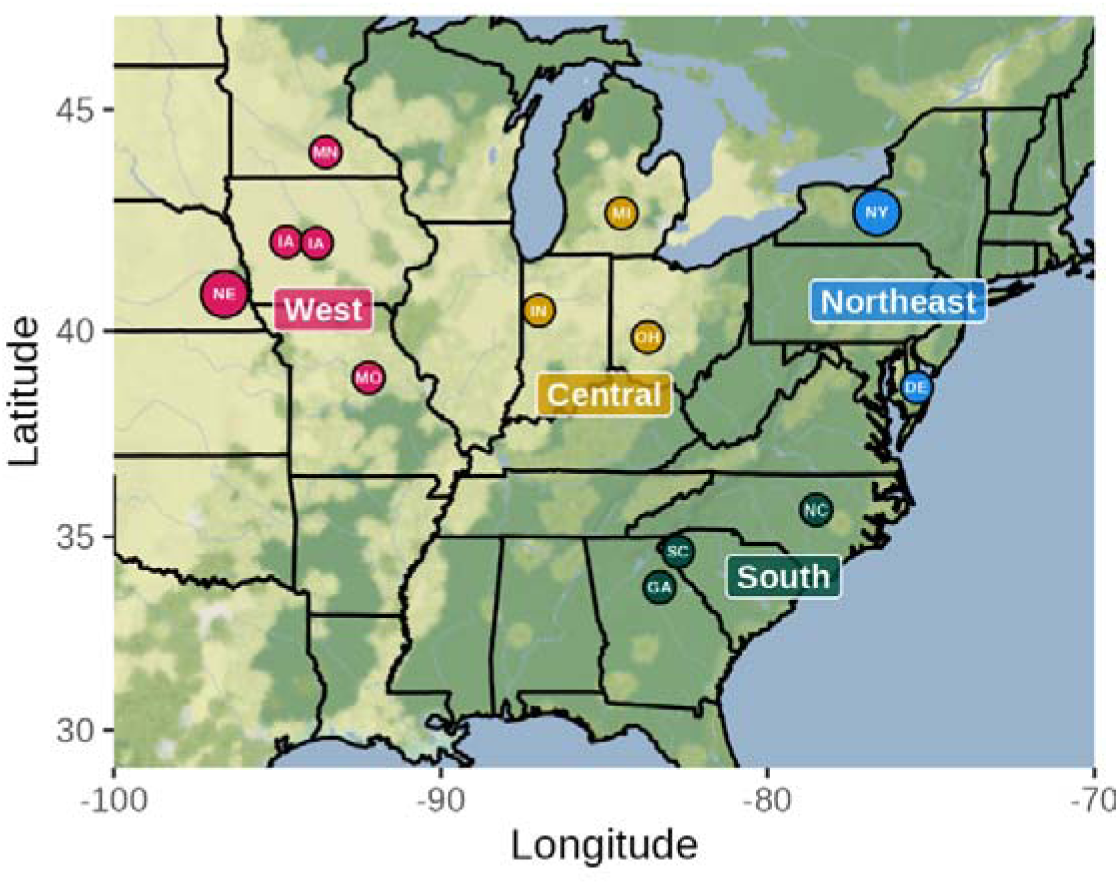
Sampling locations. Sample locations for the final, filtered dataset are shown, grouped by major geographic regions. A total of 15 fields were sampled and passed filtering; the Nebraska (NE) and New York (NY) points are larger to indicate they contain 2 sites close enough that their markers completely overlap. Map background courtesy of Stamen Maps via the ggmap package **[61]**.

### 4.1 Core bacterial community of maize stalks

The “core” microbiome refers to taxa that are present across most samples of a given dataset. There is no single accepted cutoff for a core microbiome in the literature [62], with some studies using anywhere from 5% [15] to 90% [14] of samples; we therefore classified any taxon present in at least 60% of samples to be “core.” We also performed this classification for the overall dataset, along with separate classifications for each location and genotype. Core taxa across all samples are in **Table 1**, while breakdowns by location and genotype are in **Supplementary Tables S3 & S4**.

**Table 1:** Core taxa across all 502 samples.

| Phylum | Class | Order | Family | Genus |
| --- | --- | --- | --- | --- |
| Acidobacteriota<br>(74.7%) | Vicinamibacteria<br>(64.5%) | Vicinamibacterales<br>(62.4%) |  |  |
| Actinobacteriota<br>(93.4%) | Actinobacteria<br>(89.3%) |  |  |  |
| Bacteroidota<br>(95.4%) | Bacteroidia (95.0%) | Cytophagales<br>(80.1%) | Spirosomaceae<br>(61.0%) |  |
| ” | ” | Flavobacteriales<br>(71.5%) | Weeksellaceae<br>(60.4%) |  |
| ” | ” | Sphingobacteriales<br>(68.7%) | Sphingobacteriaceae<br>(64.9%) |  |
| Chloroflexi (70.7%) |  |  |  |  |
| Firmicutes (76.3%) | Bacilli (71.7%) |  |  |  |
| Planctomycetota<br>(70.5%) | Planctomycetes<br>(64.1%) |  |  |  |
| Proteobacteria<br>(99.8%) | Alphaproteobacteria<br>(98.8%) | Rhizobiales<br>(90.0%) |  |  |
| ” | ” | Sphingomonadales<br>(77.7%) | Sphingomonadaceae<br>(77.7%) | *Sphingomonas*<br>(70.1%) |
| ” | Gammaproteobacteria (99.4%) | Burkholderiales (93.9%) | Comamonadaceae (80.9%) |  |
| ” | ” | Enterobacterales (69.1%) |  |  |
| ” | ” | Pseudomonadales (79.1%) |  |  |

Percentages indicate the percentage of samples each taxon was detected in at 0.1% abundance or greater.

The most commonly detected phyla were the Proteobacteria (nearly 100% of samples), Bacteroidota (95%), and Actinobacteria (93%). At finer taxonomic levels, the most consistently identified taxa were the Bacteroidia (95%) and Alpha- and Gammaproteobacteria classes (∼99% each), the orders Rhizobials (90%) and Burkholderiales (94%), and the families Comamonadaceae (81%) and Sphingomonadaceae (77%). Only one genus (*Sphingomonas*, 70%) was core when considering all samples. In terms of read counts, core phyla constituted an average of 89% of all reads, with lesser amounts for classes (76%), orders (52%), families (21%), and genera (4%) (**Supplemental Figure S1**). This implies that the stalk microbiome is conserved at broad taxonomic scales (phylum, class) but becomes increasingly variable at finer divisions (family, genus).

Calculating the core microbiome within each location reveals a large number of conserved taxa, which follows the same trend of more conservation at higher taxonomic levels (**Figure 2A**). Focusing on a single taxonomic level (class, **Figure 2B**) shows the trend of a few taxa being core across all or nearly all locations (Bacilli, Actinobacteria, Bacteroidia, and Alpha-and Gammaproteobacteria), whereas others are core only in a subset of locations (e.g., Anaerolineae, Verrucomicrobiae, Planctomycetes) and yet others are core only in one or two locations (e.g., Acidimicrobiia, Chloroflexia, Blastocatellia). The same pattern occurs when the core microbiome is calculated across genotypes (**Figures 2C & 2D**). An intriguing pattern are taxa that are highly prevalent in one location/genotype but rare or absent in others. For example, bacteria in the class Deinococci are common in the MNH1 location (92.3% of samples) yet rare (<10% of samples) in INH1 and NEH1 (**Figure 2B; Supplemental Table S3**). Such location-specific differences in prevalence could potentially impact plants’ interactions with the local microbiome.

**Figure 2:**
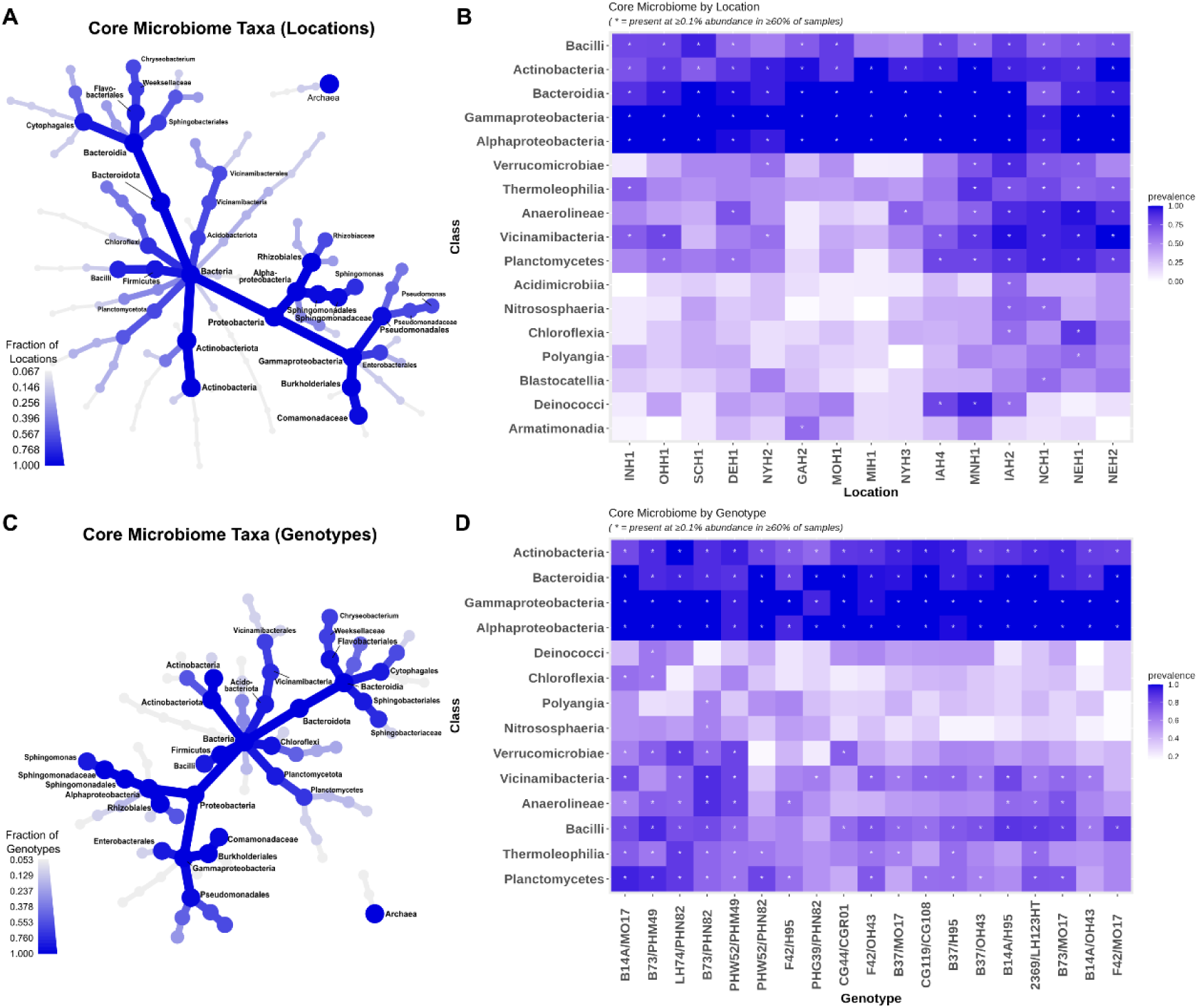
Core microbiome by location and genotype. The core microbiome was calculated separately across locations (A, B) and genotypes (C, D). At every taxonomic level (Kingdom, Phylum, Class, etc.), a taxon was considered “core” if it constituted at least 0.1% of reads in at least 60% of samples. Taxonomy trees (A, C) show all taxa that were core in at least one location/genotype; labels indicate taxa that were core in at least 60% of locations/genotypes. Heatmaps show the relative abundance of bacterial Classes (y-axis) across locations (B) and genotypes (D); both axes are ordered based on hierarchical clustering of the data, so similar taxa and locations/genotypes group together. Asterisks (*) indicate whether that Class is considered core for that location or genotype. See **Supplemental Tables S3 & S4** for the complete list of core taxa by genotype and by location.

### 4.2 Microbial Diversity Significantly Differs Based on Environment and Genotype

To further characterize the maize stalk microbiome, we calculated basic diversity metrics for all samples. Alpha diversity (microbial diversity within individual samples) was estimated for each sample using the number of observed ASVs, the Simpson index, and the Shannon diversity index. The number of observed ASVs represents the richness of the microbiome community (how many unique microbes exist in a sample), while the Simpson and Shannon indices also incorporate the community’s evenness.

All three alpha diversity metrics differed significantly among environments when tested by the Kruskal-Wallis test (p < 0.05; **Figure 3; Supplementary Table S5**). However, no significant difference was observed for any alpha diversity metrics among maize genotypes (Observed ASV: p-value: 0.40; Shannon p-value: 0.44; Simpson p-value: 0.46).

**Figure 3:**
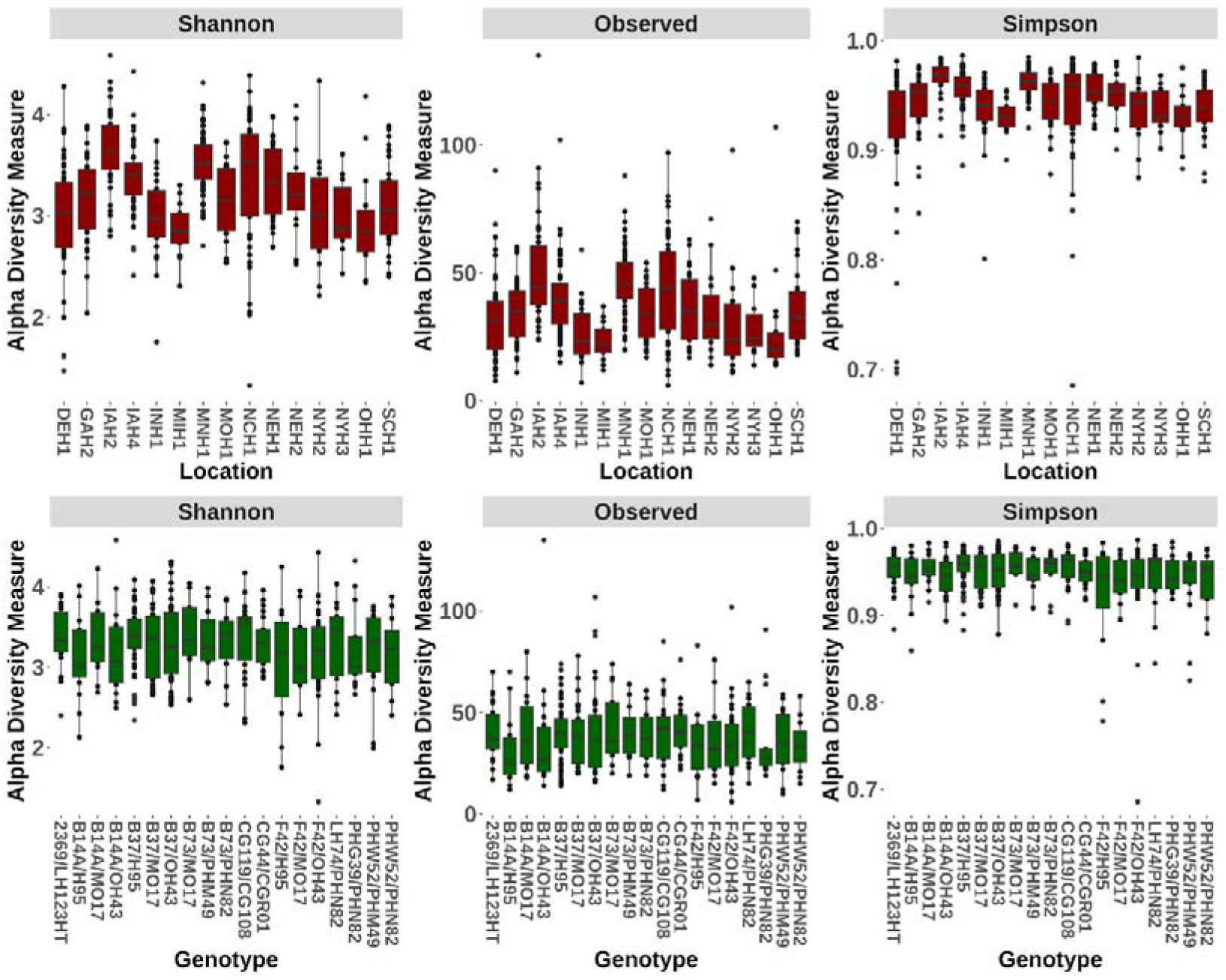
Stalk microbiome alpha diversity. Plots show the measured Shannon diversity (left), Observed ASVs (middle), and Simpson index (right), split by location (top, red) and genotype (bottom, green). In each plot, higher values indicate more diverse endophyte communities. Kruskal-Wallis tests of significance indicate that environments significantly differ from each other in alpha diversity, but genotypes do not. (See main text for details).

We also evaluated the differences in microbial communities using beta-diversity (the differences among samples), calculated using the weighted and unweighted UniFrac distances [63] (**Figure 4; Supplementary Figure S2**). We found that these distances were highly significantly different between environments, as tested by PERMANOVA, and maize genotype was also significantly associated, albeit less strongly (**Supplementary Table S6)**.

**Figure 4:**
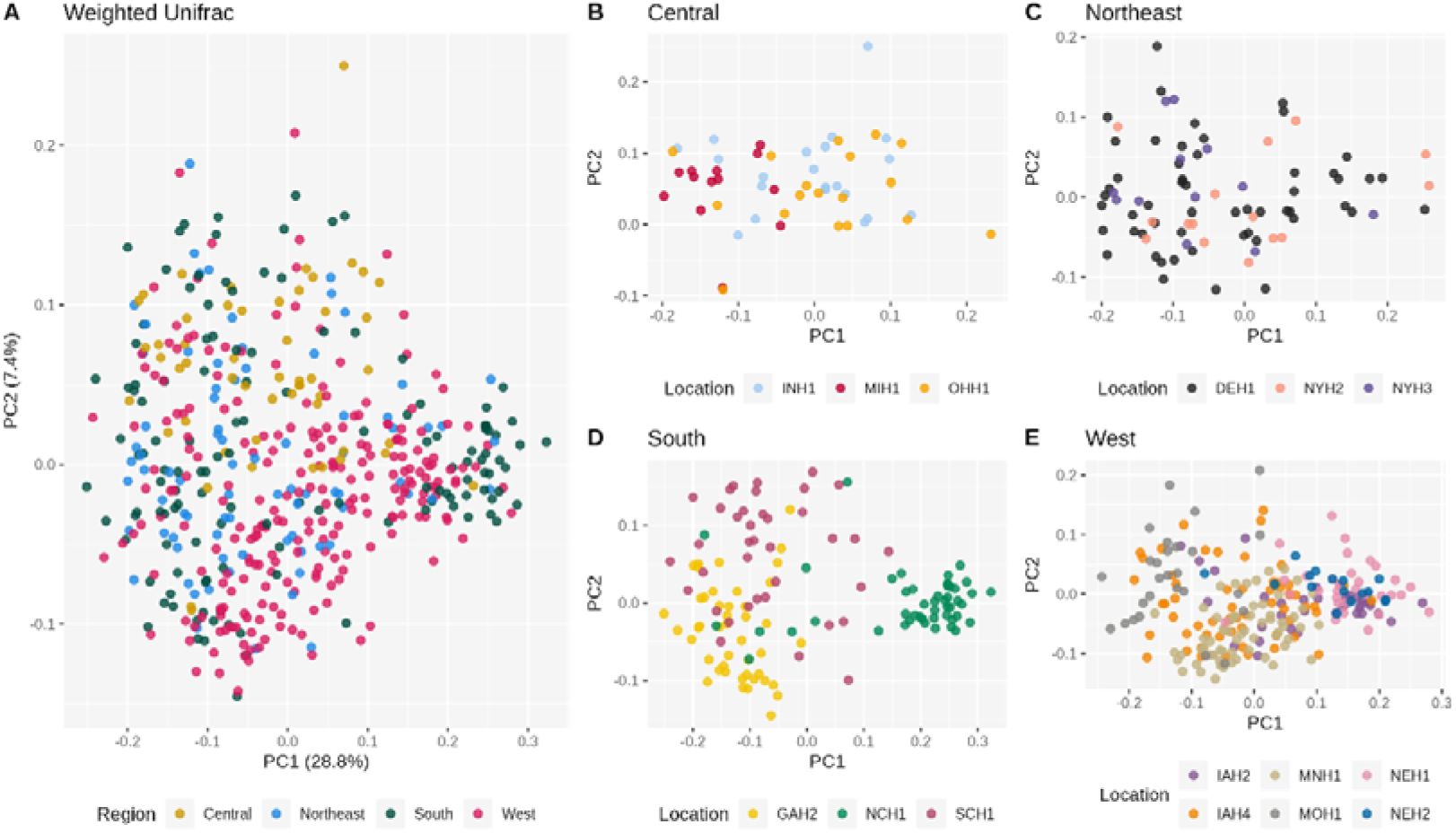
Beta diversity of the maize stalk microbiome across locations. Weighted UniFrac distances were calculated among all samples, and the first two principal coordinates plotted for all samples (A) and for each region (B-E). PERMANOVA of the total dataset indicates that samples cluster significantly based on location (p=0.0001; 9999 permutations).

### 4.3 Environment and GxE interactions are the major drivers of the maize endophyte community

A major goal of this research was to quantify the relative impact of genetics, environment, and their interaction on the maize stalk microbiome. To this end, we estimated the effect of environment (field location), maize genotype, and their interaction using linear regression models for alpha diversity and distance-based redundancy analysis (dbRDA) for beta diversity. The two biggest contributors to variance in alpha diversity were environment and GxE interaction, which explained 20-22% and 26-28% of the variation in alpha diversity metrics, respectively (**Figure 5A**). Maize genotype only explained 3-4% of the variation, similar to the non-significant results with the Kruskal-Wallis test (**Supplementary Table S7**). For beta diversity, environment, maize genotype, and GxE interaction respectively explained 12-22%, 5%, and 23-25% of the total variation for UniFrac distances; Bray-Curtis and Jaccard distances showed much smaller effect of environment (3%) and nonsignificant effects of genetics and GxE (**Figure 5B; Supplementary Table S8**).

**Figure 5:**
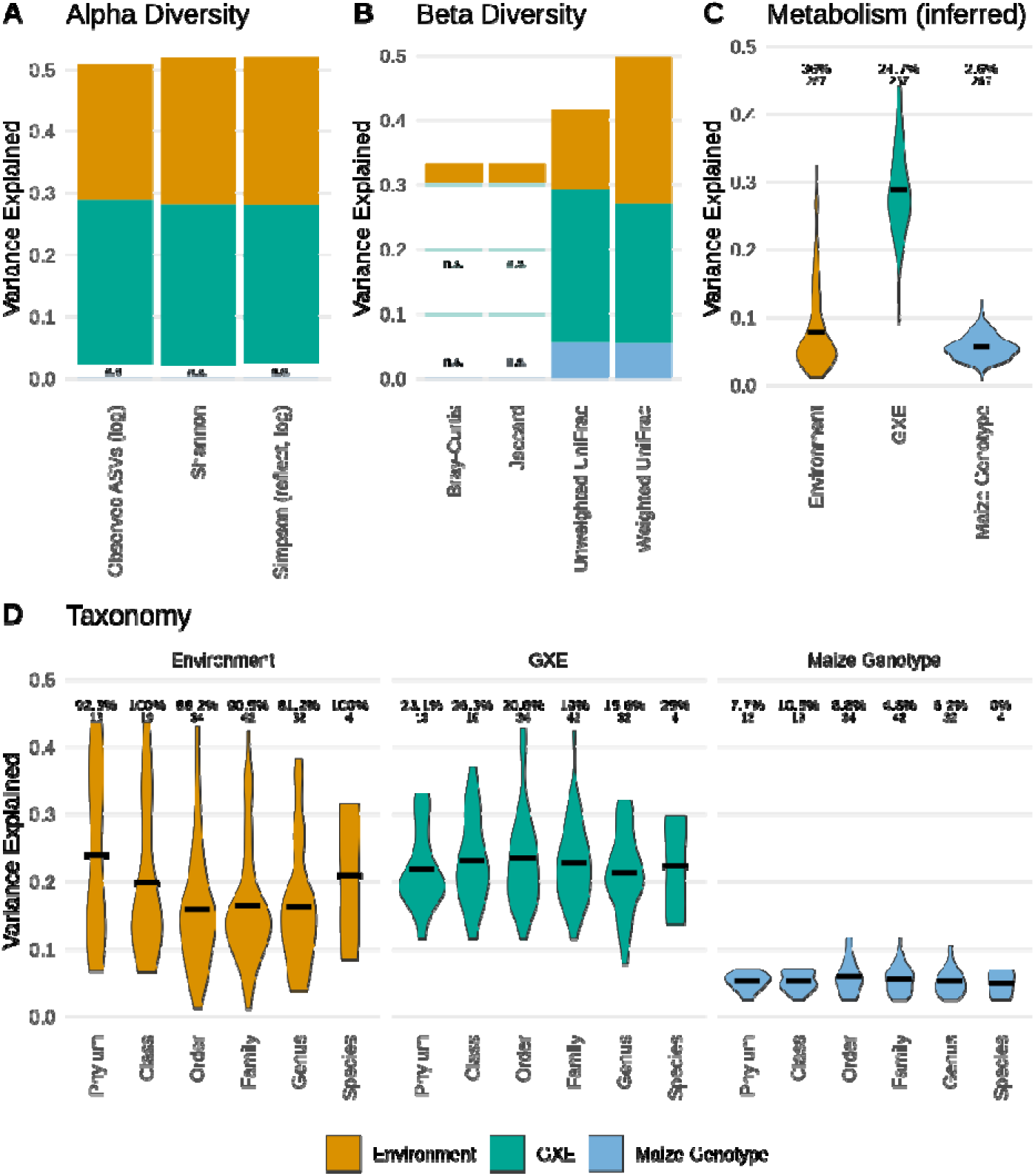
The relative impact of genetics and environment on the stalk microbiome. The proportion of variance of (A) alpha diversity, (B) beta diversity, (C) inferred metabolism, and (D) taxonomy of the stalk microbiome. In A & B, “n.s.” and transparent colors indicate non-significant terms (p>0.05). In C & D, both the total number of tests and % significant are shown. (Part C used a false discovery rate cutoff of q < 0.01 due to the large number of tests; part D used a straight p < 0.01 cutoff.) Percent variation explained and significance were calculated by linear regression (A, C, D) and distance-based redundancy (B).

When looking at microbial communities, functional capacity is often considered a better representation of community makeup than taxonomic identity. (That is, what the community can do is more important than who is present.) This is usually surveyed with shotgun metagenomics; however, there are currently no methods to reliably separate endophytes from their (much more abundant) plant host for sequencing. Instead, we used PICRUST2 [64] to estimate the predicted metagenome functionality for each sample based on its 16S rRNA gene profile. We then calculated the contribution of environment, GxE, and maize genotype to the abundance of the individual predicted metabolic pathways (**Figure 5C; Supplementary Table S9**). In this case, GxE (30% of variation) showed a significantly larger impact than either environment or genetics alone (5% each). This pattern also held when only the terms that were statistically significant in ANOVA are included in the calculation (**Supplementary Figure S3**; median 12%, 10%, and 39% for G, E, and GxE respectively).

We also investigated how environment, maize genotype, and GxE interaction contributed to the abundance of individual taxa at various taxonomic levels. Since rare taxa show an extremely high GxE effect due to being present in only a few environments, we filtered the data only to keep taxa present in at least 20% of samples and added a pseudocount of 1 to all remaining abundances. As with the diversity metrics, environment and GxE interaction were the major drivers of individual taxa, and the breakdown was consistent across different taxonomic levels (**Figure 5D; Supplementary Table S10**). Although environment explained ∼15% of the variation for most taxa, it explained 30-40% for a small subset, including Vicinamibacterales, Spirosomaceae, Deinococcaceae, and Pseudomonadaceae. The median contribution of the GxE effect on individual taxa was ∼ 23.6%, with Azospirillaceae, Moraxellaceae, and Anaerolineae being the three bacteria families with the highest impact of GxE interaction.

### 4.4 Maize endophyte communities are associated with environmental factors between experiment fields

Since the Genomes to Fields project includes environmental sampling of soil parameters at the beginning of the season, we used linear regression to relate individual environmental components to community metrics. Since each field location has only a single set of environmental parameters, metrics were calculated by either combining (beta diversity) or averaging (alpha diversity; inferred metabolism) all samples within a location.

The soil pH had a negative association with both the diversity and richness of endophyte microbiome communities. Although the direct soil pH was weakly negatively associated with diversity (**Figure 6, A-C**), the Woodruff (WDRF) buffer pH was strongly and negatively associated with diversity (**Figure 6, D-F**). While many locations share a buffer pH of 7.2, those that do also are generally the least diverse among all samples.

**Figure 6:**
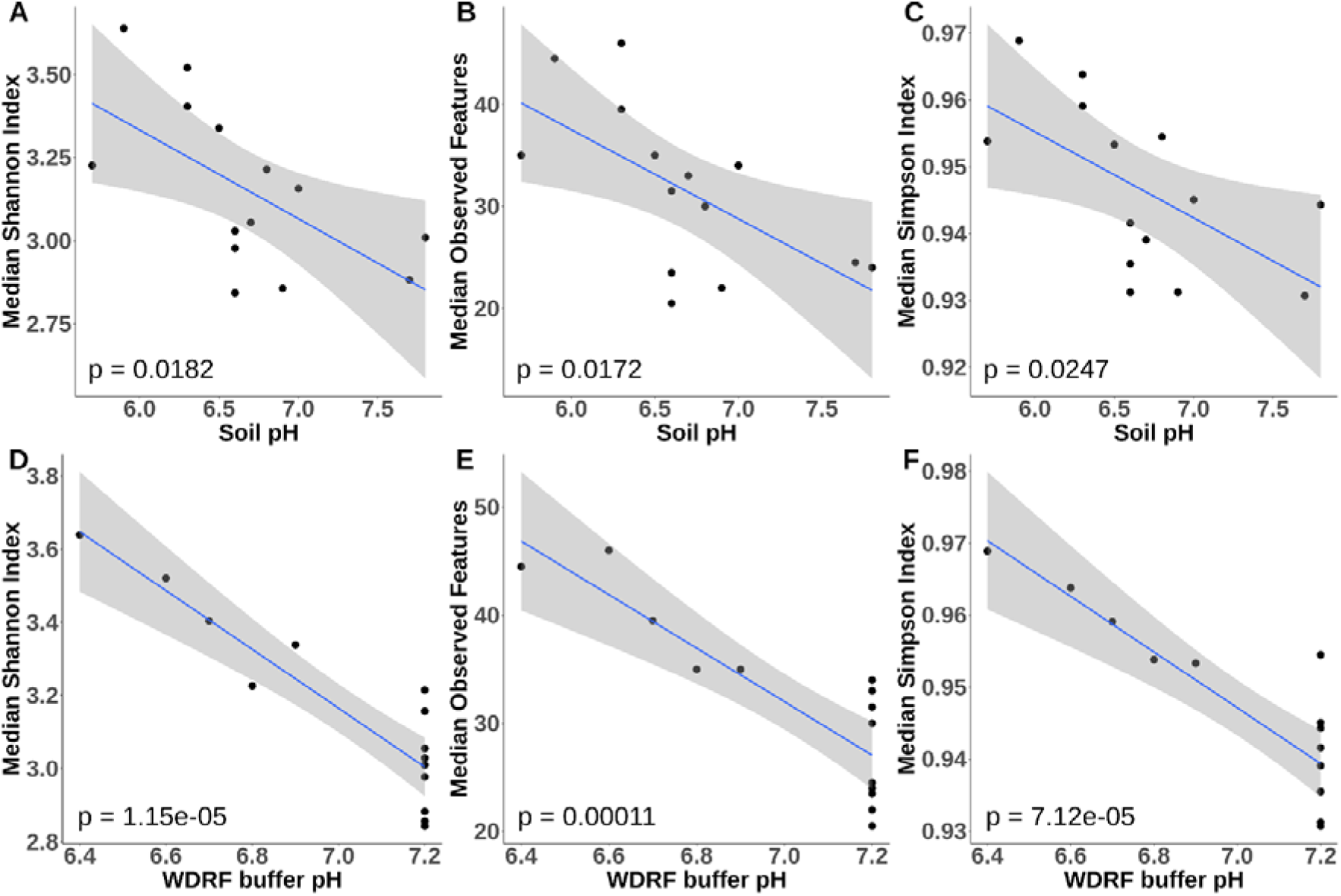
Soil pH is negatively associated with stalk bacterial diversity. (A-C) Each of the 3 different alpha diversity measures is weakly negatively associated with direct soil pH, whereas (D-F) the Woodruff (WDRF) soil buffer capacity is more strongly negatively associated.

For beta diversity, we found a significant relationship between Weighted UniFrac distances and soil potassium concentrations (r=0.65, p=0.001) (**Figure 7**).

**Figure 7:**
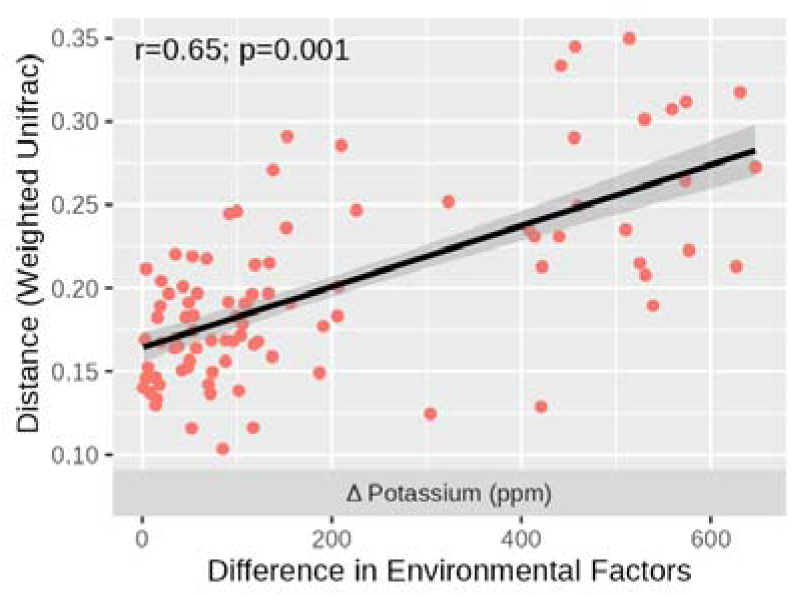
Relationship between beta diversity distances and soil potassium. Each point represents a single pairwise comparison between locations, with the difference in potassium concentration on the X axis and the Weighted UniFrac distance of the stalk microbiome on the Y axis. The correlation (test statistic) and p-value from a Mantel test with 999 permutations are shown at the top.

Finally, we tested the association between the average abundances of predicted metabolic pathways at each location and local environmental factors using linear regression. However, after multiple testing correction no pathways showed significant associations with environmental factors.

## 5 Discussion

Our results demonstrate that environmental and GxE interaction are the driving forces that shape the bacterial community living in maize stalks. This pattern held across every type of measure we tested, including individual taxa abundance, community diversity, and functional capacity. Conversely, the genotype of the maize plant by itself had little consistent effect. Similar results have been found for the maize rhizosphere [16] and switchgrass roots [14], implying this may be a general trend for plant-associated microbiomes. In contrast, an analysis of poplar root microbiomes showed a large effect for local soil and little effect of plant genotype, but no significant GxE interaction [65]. However, the smaller sample size (<30) and number of genotypes (4) may have limited power for the latter.

The idea of a nonsignificant main effect of genetics but significant GxE interaction can be difficult to conceptualize. In essence, these results mean that host genetics are a significant influence but that the effect of those genetics is almost wholly determined by the environment they are in. Another way to phrase this is that host genetics are consistent within an environment but inconsistent across environments, so knowledge of how a particular genotype shapes its microbiome in one environment gives little to no information about how it will do so in a different environment.

One implication of our results is that even though plant genetic effects have little significance on their own, they will appear significant if only tested within a single environment. When the environment is held constant, the GxE effects get folded into simple genetic effects; only when multiple environments are involved can GxE be separated out. This likely explains how many prior groups—including our own—were able to find significant genetic effects since they often limited the experiment to a single environment [66–68]

This work also adds to the understanding of plant microbiome interaction by focusing on bacterial stalk endophytes. Most of the previous publications in this area are focused on the rhizosphere [14,69–71] or phyllosphere [13]. By focusing on stalk endophytes, this study rounds out understanding the maize microbiome and confirms that similar mechanisms are at play in each of its major compartments.

### 5.1 Environmental Factors Affecting Maize Stalk Endophytes

Of the various environmental factors tested, we found that soil pH and soil potassium were the most important influences on the endophyte community. Since these factors are both soil factors, but the community we studied was in the maize stalk, it seems likely that these factors are working indirectly, either by determining the soil microbial community which can colonize the plant interior by entering through the roots [72,73] or altering the physiological state of the maize plant.

Soil pH has previously been found to be one of the largest determinants of the soil microbial community [74,75]. At least some of the maize endophyte community derives from the soil [76], though there is debate as to how much [77]. Taken together, these data imply that soil pH is most likely shaping the soil microbiome as opposed to affecting the stalk community directly.

Potassium, in contrast, is one of the most important nutrients for plant growth, and it is required by numerous plant growth processes such as enzyme activation, stomatal activity, water and nutrient transport, and photosynthesis [78]. The recommended range of Potassium for maize production is 100-200 µg g^−1^ [79], and soil test analyses indicate that locations spanned a range of levels, with 3-4 locations in each set of <100 µg g^−1^, 100-150 µg g^−1^, 150-200 µg g^−1^, and >200 µg g^−1^ at the start of the season. (Soil data at the time of sampling was not available.) Endophytes such as *Enterobacter cloacae, Bacillus pumilus,* and *Pseuodomonas sp.* are known to enhance potassium uptake by solubilizing unavailable potassium stored in the soil [80]. The inoculation of endophytic *Burkholderia* sp., meanwhile, can increase the root length and dry weight of maize seedlings by solubilizing potassium from glauconite and muscovite [81]. The increased soil potassium concentration might favor these potassium-solubilizing endophytes and thus change the endophyte community composition. Soil potassium is also a significant factor in poplar root microbiomes [82], implying it may function as a general determinant of plant-associated microbial communities.

Given the importance of nitrogen as a limiting nutrient, one might expect soil nitrogen to also be a significant factor. However, because all samples came from agricultural fields that were fertilized to standard levels for maize production, variation in soil nitrogen content was much smaller (5.8 - 54.0 µg g^−1^ nitrate versus 55 – 702 µg g^−1^ potassium, ignoring any side-dressing [generally nitrogen] added to the field after the initial soil samples). It may be that in these highly managed and fertilized fields, nitrogen levels are too similar to identify any differences due to soil nitrogen content.

### 5.2 Implications for the core maize microbiome

The “core microbiome” describes a set of microbial taxa that are consistently present across many samples. This is an intuitively useful concept that helps organize what taxa one can expect to find in a given environment. However, there is no standard way to define what constitutes a core microbe [62], and our data show that—at least for bacteria in the maize stalk—the core is highly dependent on how one looks at the data. Whether across all samples or looking within genotypes and locations, we found significant conservation at broad taxonomic scales (phylum, class) but very little at fine scales (genus, species) (**Figure 2**). Moreover, while some taxa were “core” across many locations or genotypes (e.g., Alphaproteobacteria), others were highly prevalent in a few locations but often absent from others (e.g., Deinococci) (**Figure 2B**). It may be more useful to consider the core microbiome as a spectrum, where some taxa are “more core” or “less core” than others while recognizing the variation that can occur across samples or among experiments.

## 6 Conclusions

These results have significant implications for future research. For one, the large effect of GxE on the endophyte community will likely make predictions of the endophyte community difficult. Accurate GxE prediction requires much larger datasets than prediction based on genetics or environment alone. Gathering data on this scale is difficult and expensive and was one of the main motivations for forming the Genomes to Fields Initiative. Few datasets on this scale exist, however, and that bottleneck will severely limit the ability to make accurate models. This is especially true as researchers look to apply machine learning methods to this sort of dataset [83], since these methods require very large datasets to train on and sift signal from the noise.

Another impact involves the development of biologicals (treatments based on living organisms, especially bacteria and fungi) for crop improvement. While there has been great interest in using biologicals to improve agriculture, most microbes that show early promise fail to translate in field settings [84]. At least some of this could be due to strong GxE interactions, as shown here, so that both environmental effects and plant genetics impact the final outcome. Although it is easiest to look for microbes that work consistently across many conditions, the strong environmental and GxE effects we see indicate that such broadly beneficial microbes may be very rare in native communities. Since many of these microbes are being developed as seed treatments, it may work better in the long run to select microbes that work well on specific genotypes or lineages and sell them together as a sort of “extended genome” or holobiont product. Whether this approach would actually work better than trying to find one microbe that works well across all possible genotypes remains to be seen.

Ultimately, our data show that the plant microbiome system is complex and influenced by a wide variety of factors. Fully understanding this system will require a thorough understanding of both the microbial community itself and the way it interacts with plant genetics and the environment, including how we manage that environment in a field setting. The plant-microbiome-environment interaction makes for a very complex system that will require concerted efforts at many different scales of investigation to fully understand. Although the complexity makes the problem difficult, it also means there is a great deal more to find, making the area rich for future research.

## 7 Declarations

### Ethics approval and consent to participate

Not applicable

### Consent for Publication

Not applicable

### Availability of data and materials

Illumina read data is available at the NCBI Sequence Read Archive, accession PRJNA883673. Genomes to Fields metadata and soil parameters are available on Cyverse [85] via the official Genomes to Fields website [86]. The bioinformatic code used for analysis is available via Github [87].

### Authors’ Contributions

**HL**: Methodology, Software, Formal analysis, Investigation, Data Curation, Writing – Original Draft, Visualization. **HKG**: Methodology, Investigation, Project administration. **JE**, **SFG**, **CH**, **JBH**, **JEK**, **NdL**, **MPS**, **JCS**, **RSS**, **SCM, PT**, and **AT**: Resources, Supervision, Funding acquisition, **DCL**: Project administration. **JGW**: Conceptualization, Methodology, Software, Formal analysis, Investigation, Data curation, Visualization, Supervision, Funding acquisition. **All authors**: Writing – Review & Editing.

## Acknowledgements

In addition to the coauthors of this paper, the Genomes to Fields 2019 experiment was also conducted by Edward Buckler and Shawn Kaeppler. Technical support was provided by Colby Bass, Jim Elder, Amanda Gilbert, Susan Melia-Hancock, Christine Smith, Nicholas Lepak, Trevor Perla, Naomi Rodman, and Bill Widdicombe. Curation of the 2019 data was done by Alejandro Castro Aviles and Dayane Cristina Lima. Original germplasm was maintained by the USDA germplasm repository. Mention of trade names or commercial products in this publication is solely for the purpose of providing specific information and does not imply recommendation or endorsement by the U.S. Department of Agriculture. USDA is an equal-opportunity provider and Employer.

## Additional Files

Additional File 1 - Supplemental Figures.docx: Supplemental Figures S1-S3. Additional File 2 – Supplemental Tables.xlsx: Supplemental Tables S1-S10.

