## Additional File 1 for "Gene-by-Environment Interaction Significantly Drives Bacterial Endophyte Communities in Maize Stalks"

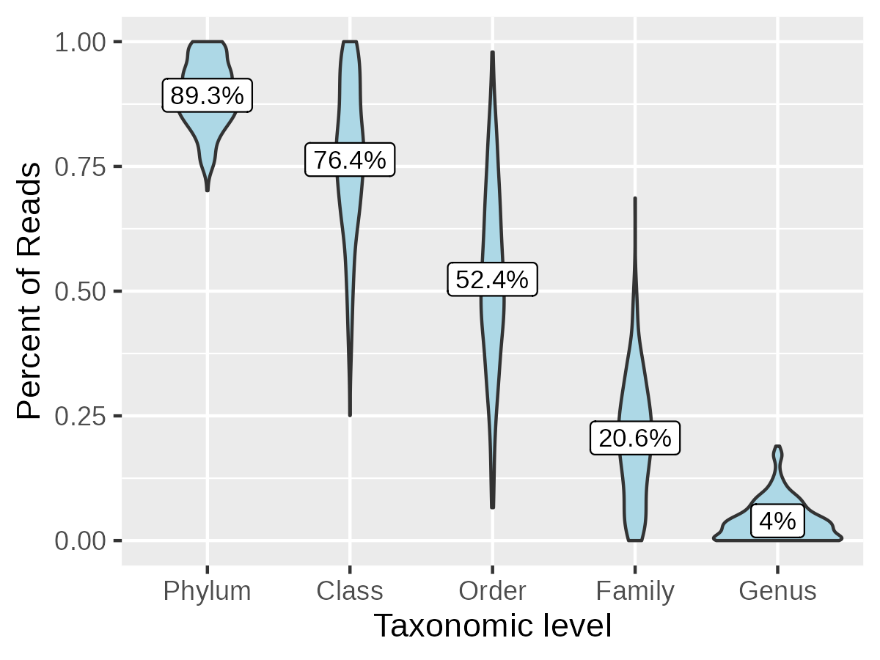

Figure S1: Percentage of the stalk microbiome belonging to “core” taxa, meaning present in 60% of samples at a minimum of 0.1% abundance. At the broad scale (Phylum, Class), most of the community belongs to core taxa, but at finer level scales the values fall off, indicating higher variability among samples.

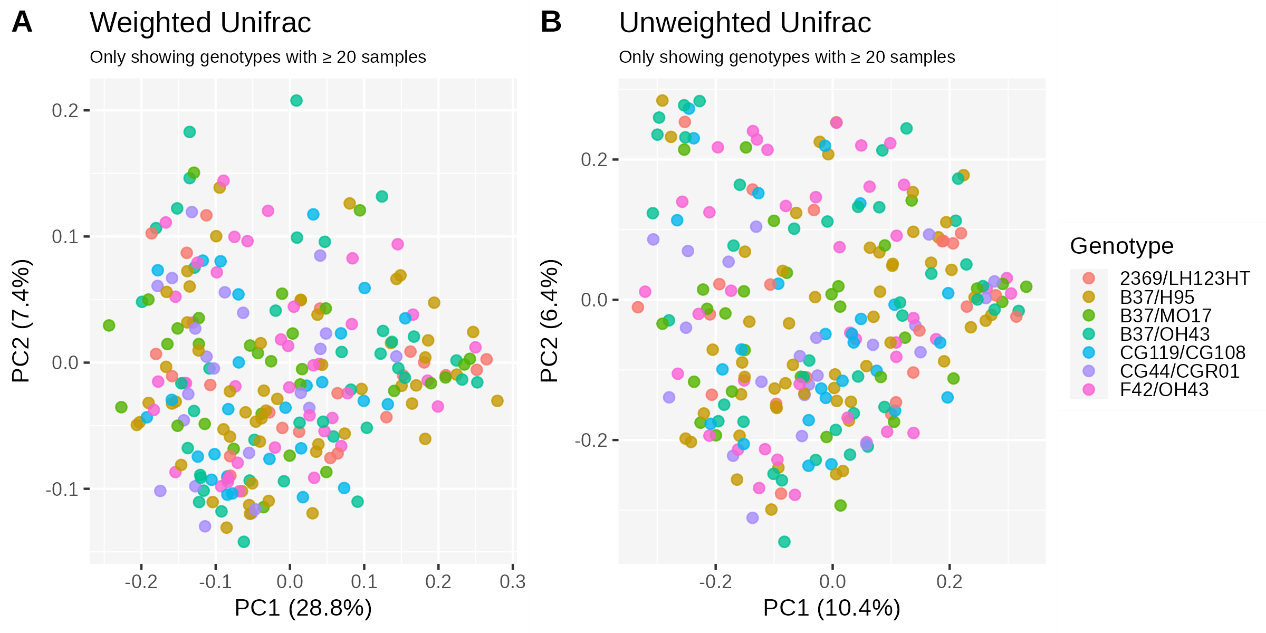

Figure S2 - PCOA of Unifrac distances among samples, colored by genotype. Both weighted (A) and unweighted (B) UniFrac distances were calculated among all samples, and the first two principal coordinates plotted, colored by maize genotype. (For simplicity, only genotypes with at least 20 samples are shown). PERMANOVA with 9999 permutations among all samples indicates that samples cluster significantly based on genotype (p=0.0033 for weighted and p=0.001 for unweighted).

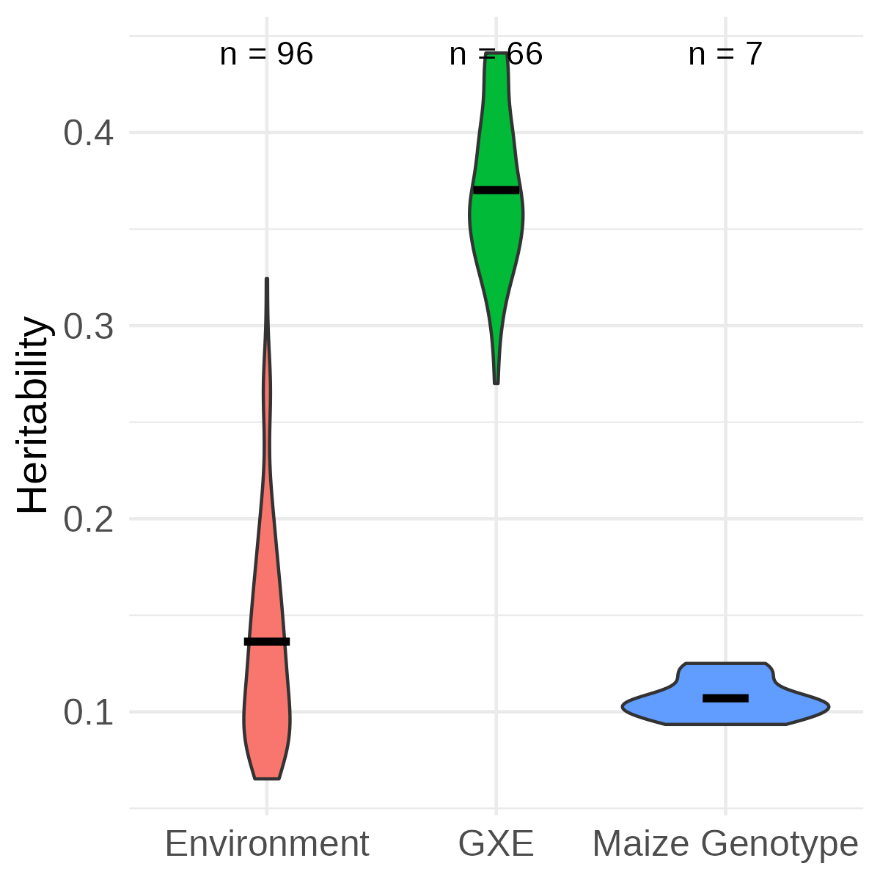

FIgure S3 - Distribution of fraction of variation explained by genetics, environment, and GxE for only statistically significant terms. The dataset is the same as Figure 5C in the main text, except only terms that showed statistical significance (FDR <= 0.01) are included in the distribution. (The number of such terms is shown above each distribution).
